# Effect of Cytoplasmic Viscosity on Red Blood Cell Migration in Small Arteriole-level Confinements

**DOI:** 10.1101/572933

**Authors:** Amir Saadat, Christopher J. Guido, Eric S. G. Shaqfeh

## Abstract

The dynamics of red blood cells in small arterioles are important as these dynamics affect many physiological processes such as hemostasis and thrombosis. However, studying red blood cell flows via computer simulations is challenging due to the complex shapes and the non-trivial viscosity contrast of a red blood cell. To date, little progress has been made studying small arteriole flows (20-40*μ*m) with a hematocrit (red blood cell volume fraction) of 10-20% and a physiological viscosity contrast. In this work, we present the results of large-scale simulations that show how the channel size, viscosity contrast of the red blood cells, and hematocrit affect cell distributions and the cell-free layer in these systems. We utilize a massively-parallel immersed boundary code coupled to a finite volume solver to capture the particle resolved physics. We show that channel size qualitatively changes how the cells distribute in the channel. Our results also indicate that at a hematocrit of 10% that the viscosity contrast is not negligible when calculating the cell free layer thickness. We explain this result by comparing lift and collision trajectories of cells at different viscosity contrasts.

## I. INTRODUCTION

Microcirculation in vascular systems is a vital component of living organisms where oxygenated blood passes through terminal arterioles (smaller blood vessels with 20 to 50 micron diameter) and capillaries (~10 microns) [15, 36]. Whole blood is a complex fluid composed of plasma as the suspending medium and three main cellular components: red blood cells (RBCs), platelets and white blood cells. Red blood cells take up to 45% volume fraction (Hematocrit) in the whole blood and are the oxygen carriers with biconcave shape with equivalent radius of about 2.82 *μ*m. Their reduced volume is about 0.65 (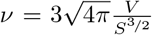, where *V* is the cell volume and *S* is its surface area) [7, 10, 26] and are highly flexible [6]. This flexibility, and resulting lift force causes the cells to migrate away from the vessel walls when they are exposed to the blood stream.

The migration of RBCs is of direct biological importance – as the cells move towards the center-line in pressure driven flows, a cell-free layer (CFL) is formed near the vessel walls (also known as Fahraeus-Lindqvist layer) [16]. This layer is known to contribute in the reduction of the blood viscosity or plasma skimming as blood perfuses the smaller vessels [13, 28]. In addition, the migration of the red blood cells induces the margination of stiffer platelets [1, 11, 19, 33, 40, 42]. Therefore, the local hematocrit of RBCs normal to the vessel walls affects the margination of platelets and their reactions with the endothelial cells in the event of an injury when the clotting cascade begins [17, 22].

Individual cell migration is partly due to a wall-induced lift force, with the velocity proportional to the stresslet, i.e., the symmetric part of the first moment of particle surface traction and inversely proportional to the squared vertical distance from the walls [39, 46]. There is another lift mechanism in pressure driven flows, due to the non-zero curvature of the flow field which drives the cells closer to the center-line, in order to minimize particle deformation [3, 4]. It has been shown that the lift velocity generally depends on capillary number Ca (the ratio of viscous forces to the membrane shear elastic forces), the viscosity contrast (λ), and the height of the channel [30].

The shape and the mode of particle rotation, namely tank-treading, tumbling, swinging, or breathing motion [2, 9, 43], are correlated with the lift velocity. More symmetric shapes and tumbling motion are expected to reduce the lift velocity. For instance, at Ca = 0.25, an RBC is in the tumbling regime [38] and therefore the migration is slower, whereas for Ca≥ 0.5, the RBC is in the tank-treading regime and lifts more quickly [30, 35]. Slipper-like and parachute-like shapes for RBCs have been extensively reported in the literature [9, 18, 20, 32, 41]. However, experiments of Lanotte et. al [21] clearly revealed existence of more complicated multi-lobe shapes. Remarkably, there is a rich spectrum of shape states and motion that RBCs may undergo depending on the initial condition [18], and Ca [38], and also the confinement level of the vessel [34].

The collective behavior of a RBC suspension is, of course, more complicated than single cell dynamics. The RBC volume fraction is 45% in whole blood, however, the physiological hematocrit levels of the micro-circulation are much lower. This reduction occurs when the blood flows from the arteries to arterioles and capillaries. The hematocrit in small vessels may be 10% or lower due to this Zweifach-Fung effect [15, 28, 37]. At these volume fractions, it is crucial to capture the interaction of the cells to describe the overall collective dynamics of an RBC suspension [23, 27, 30, 31]. In fact, the average local hematocrit of RBCs has a nontrivial form in microchannels that is a strong function of the channel size and is a direct result of the interplay between the hydrodynamic lift and cellular interaction [29]. Collision models have been developed over the last decade to approximate the form of the concentration distribution and a simple advection-diffusion equation with only binary diffusion can be successful in predicting these distributions for relatively wide channels [27, 29].

Viscosity contrast affects the shape, mode of motion, and collision dynamics of RBC suspensions [5, 14, 21, 25, 38, 44, 45]. A large body of simulations in the literature have been conducted with the viscosity ratio in the range of λ ≤ 3 [8, 12, 23, 30, 46], however, the physiological value is nearly 5 [21, 36]. For boundary integral simulations, high viscosity ratio may result in an undesirable mode of rotation [23], buckling [12] or a numerical instability [24, 25, 29]. The literature regarding the conditions under which the viscosity ratio affects RBC suspension behavior is small and incomplete. 2D LB simulations of [37] in a bifurcated channel suggest that there is a significant impact on the local RBC distribution of the parent channel when λ = 10 compared to λ = 1. Recent LB simulations of de Haan et al. [5] indicate an increasing impact of the viscosity contrast for higher flow strength (Reynolds number). In this study, simulations were performed for large channels (70 *μ*m in diameter) and the researchers observed negligible importance of λ on the cell-free-layer and also on the local hematocrit for Ht values down to 20%.

In this work, we utilize a finite volume flow solver and finite elements to model the physics of RBCs inside channels whose dimensions are ≤ 50*μ*m to represent the confinement-level similar to the small micro-circulation. We investigate the role that channel size has on the RBC distribution and the time dependence of its evolution. We show the effect of viscosity contrast (λ) on the cell distributions in these small channels at physiological hematocrit 10-20%. We report that in these smaller channels, the viscosity contrast has a measurable impact on the CFL thickness if the concentration of RBCs is at the low end of the physiological range and explain these results based on the measured lift and binary collisions observed for single cells and cell pairs.

## II. METHODS

### A. Governing Equations and Methodology

We consider the dynamic problem of an elastic RBC suspended in an incompressible Newtonian fluid, where the suspended RBC is neutrally buoyant. The total domain under consideration is defined to be Ω which will be broken into two sub-domains Ω^f^ and Ω^m^ which represent the volume of the fluid and the suspended RBC respectively (the m superscript given here to represent “membrane” as opposed to “fluid”). The governing equations are conservation of momentum in both the fluid and membrane sub-domains as well as continuity:

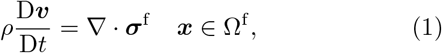

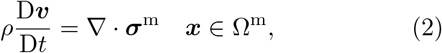

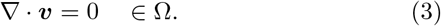

We have defined the stress in the RBC and fluid to be ***σ***^m^ and ***σ***^f^ respectively. At the boundary of contact between the membrane and the liquid we also require a stress balance to be satisfied. We denote the boundary with an outwardly-pointing unit normal ***n***. We write the stress balance condition as

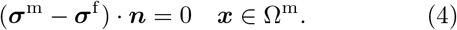

We represent the suspending fluid stress as a Newtonian stress,

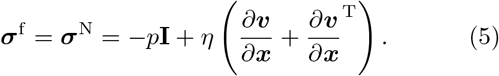

In the above, we have defined *p* to be the hydrodynamic pressure and *η* to be the Newtonian fluid viscosity.

We also must specify a constitutive model for the RBCs. The RBC will be modeled as a 2D hyper-elastic membrane. The second Piola-Kirchhoff tension, **Ŝ**, is calculated using the principle of virtual work from the strain energy areal density 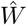, which is a function of the invariants of the right Cauchy-Green tensor 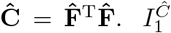 and 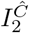 are the two independent invariants of **Ĉ** in this reduced system and the following relationship is utilized to calculate stress:

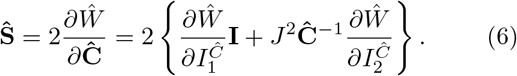

The well-known Skalak model is used for the energy areal density:

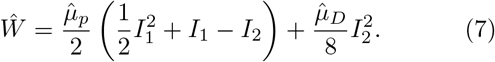

where 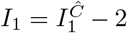 and 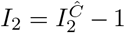 are the two invariants of the Skalak model. The Skalak model is generally used to enforce local area-incompressibility in a membrane so the dilatational modulus, 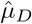, is set to be much larger than the shear modulus, 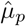.

Since we have neglected the out of plane forces in the membrane approximation, we also include the bending energy in our model. This provides an additional energy density function for bending:

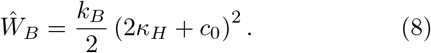

In the above expression we have defined *k_b_* as the bending modulus, *c*_0_ as the spontaneous curvature of the membrane, and *κ_H_* as the mean curvature of the membrane. consider To solve the coupled fluid-solid problem we utilize an Immersed Finite Element Method (IFEM). More details about this method can be found in a recent publication and in the supplemental information [35].

### B. Non-dimensional Equations

We can write all of the governing equations in a standard non-dimensional form. We will consider the case of Poiseuille flow in this numerical study, but without loss of generality we can choose our characteristic length to be that of the particle *R*\* (the reduced radius, about 2.82*μ*m for a RBC), the characteristic velocity to be 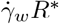 (the wall shear rate times the particle size), the characteristic stress in the fluid to be the 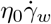, the characteristic stress in the particle to be 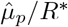, and the timescale to be 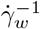. This gives us the following non-dimensional equations to solve (where non-dimensional variables and operators are indicated with a star):

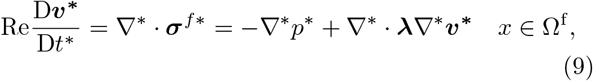

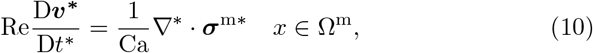

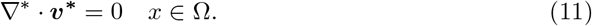

We note that the viscosity may not be constant everywhere in the simulation. There are actually two different zero shear viscosities inside and outside the membrane which we will call: *η*_in_ and *η*_out_ = *η*_0_. Additionally, non-dimensional relationships for the energy areal density are as follows for the Skalak Model and the bending energy:

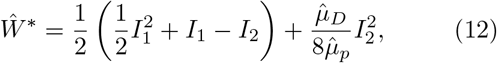

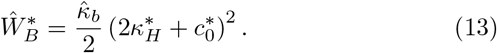

This leaves us with a total of 6 dimensionless parameters: The Reynolds number 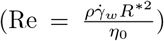, the capsule viscosity ratio 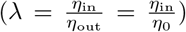, and the capillary number 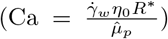 appear in the evolution equations. Additionally, two ratios appear in the constitutive equations for the solids: 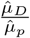 and 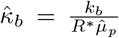. For the studies presented in this paper the Reynolds number will be smaller than 10^−1^, and the capillary number will be set to a value of 1 for all simulations. Since we desire a simulation of RBCs using the Skalak model and these systems are largely surface incompressible we will set the dimensionless ratio 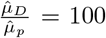. The bending parameter, 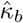, is generally much smaller than 1 [38, 46] and we set it to be 0.03 for all of the simulations. The last dimensionless parameter is a geometric parameter which is the confinement ratio of our channel, 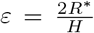. Our main aim in this manuscript is to discuss the effects that varying viscosity ratio, 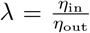 has on the cell dynamics of single and multiparticle red blood cell suspensions. Additionally we will consider the effect of using two distinct channel heights (two values of *ε* = 0.28, 0.17 corresponding to 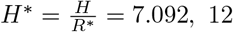).

## III. RESULTS AND DISCUSSION

### A. Single Cell and Binary Collision Behavior

We first consider the effect that the viscosity ratio (λ) has on the dynamics of single cells or of pairs of cells. Often, as mentioned above, for reasons of computational costs the viscosity ratio is assumed to be 1, but a realistic RBC has a viscosity ratio much closer to 5. In the following section we consider how the viscosity ratio affects the hydrodynamic lift of cells and also how the binary collision is affected by the viscosity ratio. Both of these features are critical since the binary collision and lift are the two main driving forces that cause cells migration. These two simple simulations can then be used as inputs into a coarse grained model such as those developed elsewhere[30].

#### 1. Hydrodynamic Lift of Single Cells

We begin by considering the behavior of single cells lifting in a channel flow for two different values of the viscosity ratio. The geometry utilized for this experiment is characterized by *ε* = 0.28 (corresponding to *H* = 20*μ*m) where in the x and z directions, we imposed a periodic boundary condition. The Ca number for these simulations is set to 1 (very close to the physiological value in micro-capillaries). Presented in Fig. 1 we see the trajectory of two cells with a viscosity ratio of 1 and 5 plotted as a function of dimensionless time 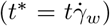. The higher viscosity ratio of 5 is plotted as diamonds and we see that the lift is substantially reduced compared to the lower viscosity ratio of 1.

**FIG. 1.**
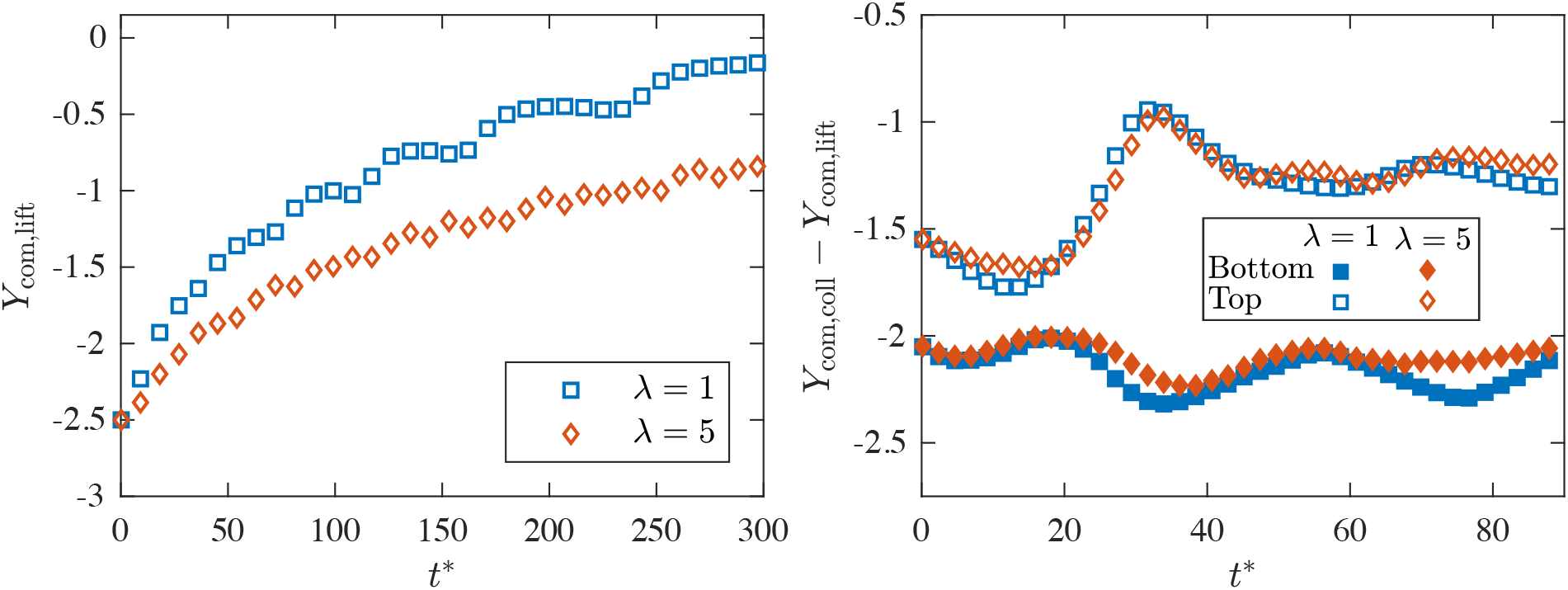
The cell trajectory as a function of time for two cells that begin close to the wall at two viscosity ratios of 1 and 5 in the left panel. The wall is located at *Y* = −3 and the time is made dimensionless with the inverse shear rate at the wall. In the right panel we plot the trajectory of two cells undergoing a binary collisions for the same viscosity ratios of 1 and 5 (squares and diamonds) where the lift trajectories have been subtracted from the motion.

This result has considerable consequences for multiparticle systems since one of the primary driving forces for RBCs to migrate toward the center of the channel is the lift [30]. The platelets are then squeezed to the perimeter by binary diffusion which leads ultimately to platelet margination and the formation of a Cell Free Layer (CFL). This result suggests that there is a smaller driving force to generate the cell free layer, since hydrodynamic lift at higher viscosity ratios is reduced.

#### 2. Binary Cell Diffusion

Next we consider the binary collision problem in channel flow. We again utilize a similar geometry used in the hydrodynamic lift study (*ε* = 0.28). The Ca is fixed to 1. The cells are initially started at fixed 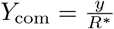 positions of −1.5 and −2 and then allowed to collide. However, the cells that collide with one another are also constantly lifting as well. We would like to remove this factor from our analysis. In Fig. 1 on the right panel we see the trajectories from this collision for both cells at two different viscosity ratios of 1 (squares) and 5 (diamonds) with the lift trajectories subtracted from the binary collision. We see the final displacement due to the binary collision has very little dependence on the viscosity ratio. This implies that the binary diffusivity for both of these systems is virtually the same. Since the lift is markedly different this allows us to conclude that the driving force for motion towards the centerline is weaker for viscosity ratio 5, but the driving force for motion away from the center is about the same (since this is binary diffusion driven [30]). In multi-particle simulations we therefore expect the CFL to be smaller for the viscosity ratio 5 systems.

### B. RBC Suspensions with 10-20% Ht

In this section, we discuss particle resolved simulations of suspensions of RBCs considering three main parameters of interest. We will consider the effect of the channel confinement (*ε*) on the distribution of cells and the dynamics of these distributions. Then we will discuss the effect of viscosity ratio (λ) and volume fraction (also known as hematocrit for RBCs which we denote as Ht) on the distribution of cells, the CFL, as well as the observed cell shapes in these flows. The simulation of the CFL size in particular is of great interest at viscosity ratios greater than 1 since to date this has not been studied for small arterioles at physiological hematocrits in such simulations (and often times it is assumed that a viscosity ratio of unity is sufficient for such calculations).

#### 1. Transient Shapes as a Function of Viscosity Ratio

We next present the effects of viscosity ratio on the shapes of RBCs in suspensions of RBCs at different hematocrit levels. Again we remark that it is known that small vessels tend to have hematocrits around 10% [15]. The first thing we examine is how the shapes of these cells differ in the two situations. It has been observed experimentally and through simulations by Lanotte et al. [21] that there are a series of unusual shapes exhibited by cells with a viscosity ratio of 5 (the physiological viscosity ratio) at relatively high shear rates. These interesting shapes include tri-lobe and multi-lobe shapes. In Fig. 2 we present snapshots of a series of shapes for different red blood cells at two viscosity ratios of 1 and 5. These cells are from a suspension simulation in a pressure driven flow with *ε* = 0.28, Ht = 10%, and at Ca = 1.

**FIG. 2.**
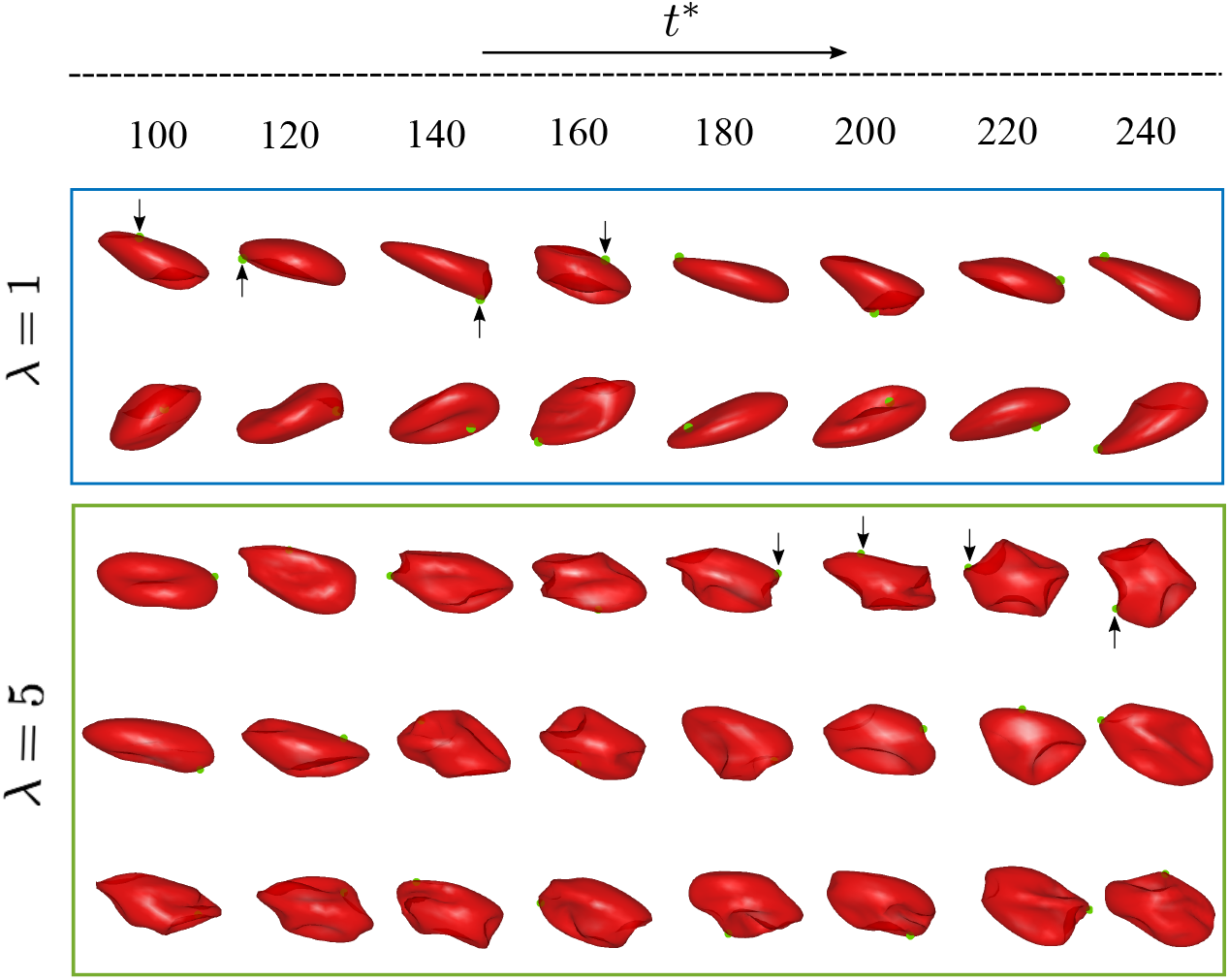
The shapes as a function of time for 5 cells from simulations with a volume fraction of 10% and channel constriction of *ε* = 0.28. The first two rows of cell images are taken from a simulation at λ =1 and we see generally smooth shapes that tank tread. The lower three rows are from simulations at λ = 5 and we see complex multi-lobe shapes with tumbling dynamics.

In Fig. 2 in the upper panel we see two red blood cells that are observed over many dimensionless times 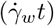. It is clear that these cells take on relatively smooth shapes such as “slippers” and are largely tank treading at this Ca number. In the lower panel we have selected three cells from a viscosity ratio 5 suspension and we observe qualitatively different cell shapes. These shapes can all be described as multi-lobe shapes such as those observed by Lanotte et al. [21]. It is clear that the viscosity ratio of 5 is required to see such shapes in our simulated system. Another clear feature is that all of these cells largely tumble.

These shapes at larger viscosity ratios that exhibit tumbling dynamics are likely important when considering the lift dynamics of individual cells. As we saw in Fig. 1, tumbling cells at the higher viscosity lift more slowly than the tank-treading lower viscosity ratio counterparts. The shape of a cell and its mode of motion strongly influence the lift of a given cell which means we expect suspensions of these cells to have different cell free layer (CFL) thicknesses and dynamics.

#### 2. RBC Distributions as a Function of Hematocrit and Viscosity Ratio

We next consider how the distributions of cells are affected by the viscosity ratio in pressure driven flow. In addition we will consider the effects of cell volume fraction Ht = 10% or 20%. Given results demonstrated for single particles and binary collision we know the fundamental driving force of lift is different between these two viscosity ratios but the binary diffusion induced by collision is largely constant. We therefore expect that the CFL and the dynamics that lead to a steady state to be somewhat different in these cells suspensions.

In Fig. 3, we see the concentration distributions at a variety of different confinements, Ht, and viscosity ratios. The top row of images are all presented at a viscosity ratio of 1, the lower row of images are simulations with a viscosity ratio of 5. If we compare Fig. 3a and Fig. 3d we see the effect of viscosity ratio at a fixed channel size of *ε* = 0.28 and fixed Ht of 10%. We see the main effect is indeed that the cell free layer has been reduced in size due to the lowered lift driving force. You can observe this reduced cell free layer by noting the depleted region near the wall on the far right of each distribution.

**FIG. 3.**
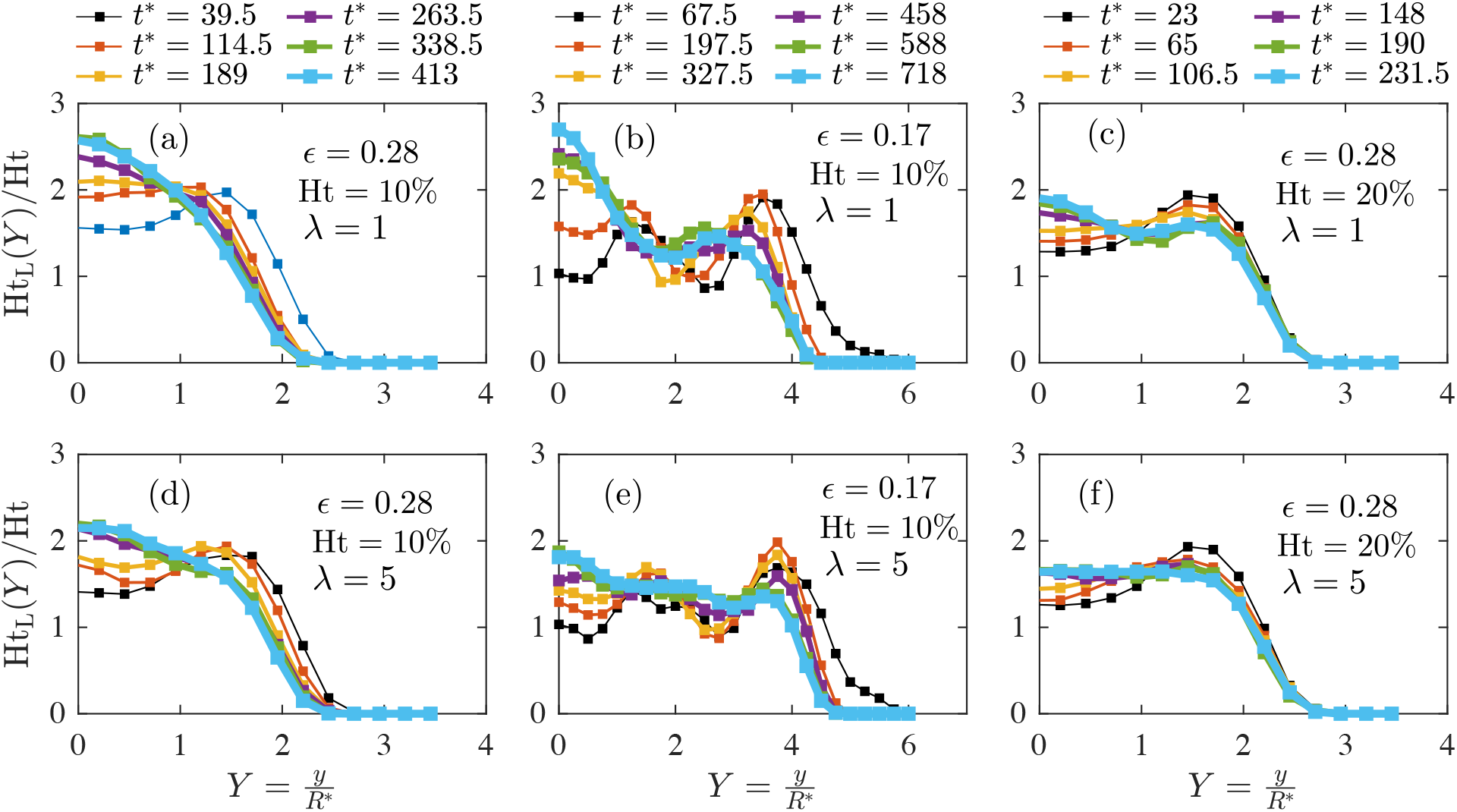
Local hematocrit in a pressure driven channel flow. The top row of images are all presented at a viscosity ratio of 1, the lower row of images are simulations with a viscosity ratio of 5. In panels (a) and (d) we consider the effect of viscosity ratio in a simulation with a fixed Ht of 10% and confinement of *ε* = 0.28. In panels (b) and (e) we consider the effect of viscosity ratio in a simulation with a fixed Ht of 10% and confinement of *ε* = 0.17. In panels (b) and (e) we consider the effect of viscosity ratio in a simulation with a fixed Ht of 20% and confinement of *ε* = 0.28.

The distribution at higher viscosity ratio has also become substantially more blunt for viscosity ratio of 5.

Additionally, we explore the channel size effects on CFL size and cell distribution for larger channel sizes with a size of *ε* = 0.17. The simulations here are conducted at a Ht = 10% and we again consider viscosity ratios of 1 and 5. In Fig. 3b and Fig. 3e the cell distributions for viscosity ratio of 1 and 5 are presented and we see similar trends as those seen at the higher confinement earlier in Fig. 3a/d. The CFL is smaller for the higher viscosity simulations and the bulk profile has become substantially more blunted. In supplementary information, we show that the second moment of the concentration profile for λ = 5 is changed by 30% compared to λ = 1.

The other feature concerning the effect of channel size can be seen from Fig. 3a/b. The smaller channel (*ε* = 0.28) on the left only shows one or two maxima depending on the time. This is very different from the results displayed in the panel (b) where there are two secondary peaks near the boundary of the CFL. These secondary peaks also seem to display similar evolution in size and position when compared to the bimodal distribution seen in the smaller channel. This evolving bi-modality of the smaller channel is very interesting to observe since smaller channels have been under-explored in this context despite their physiological relevance. This feature in the cell distribution undoubtedly has implications in how cells split and marginate in junctions with varying inlet sizes which is likely critical in physiological conditions.

In Fig. 3c and Fig. 3f we consider the same comparison but at the higher volume fraction of Ht = 20%. This set of simulations were also conducted at Ca = 1 and a confinement of *ε* = 0.28. The features presented in the set of simulations are quite different. At the higher volume fraction of 20% the distributions appear to be very similar. At high concentrations the cell free layer appears to be controlled by the finite volume the cells can occupy and/or binary collision, instead of the lift that were previously much more important in lower volume fraction simulations.

We can quantify the CFL from these simulations and examine the effects of confinement, Ht, and viscosity contrast. In Fig. 4 we see the CFL plotted as a function of time for the simulations presented in Fig. 3. The steady results from the collision theory by [29] are presented as dark dashed lines. In Fig. 4a we see the CFL plotted for the four simulations presented in Fig. 3a/c/d/f as a function of dimensionless time. The higher viscosity ratios of 5 are plotted as diamonds and the lower viscosity ratios of 1 are plotted as squares. Unsurprisingly, at the higher volume fraction the CFL is much smaller (due to the excluded volume and much higher cell load). We see that the observation we made about the CFL in Fig. 3c/f is true quantitatively: the CFL is virtually independent of viscosity ratio at the higher volume fraction of 20%.

**FIG. 4.**
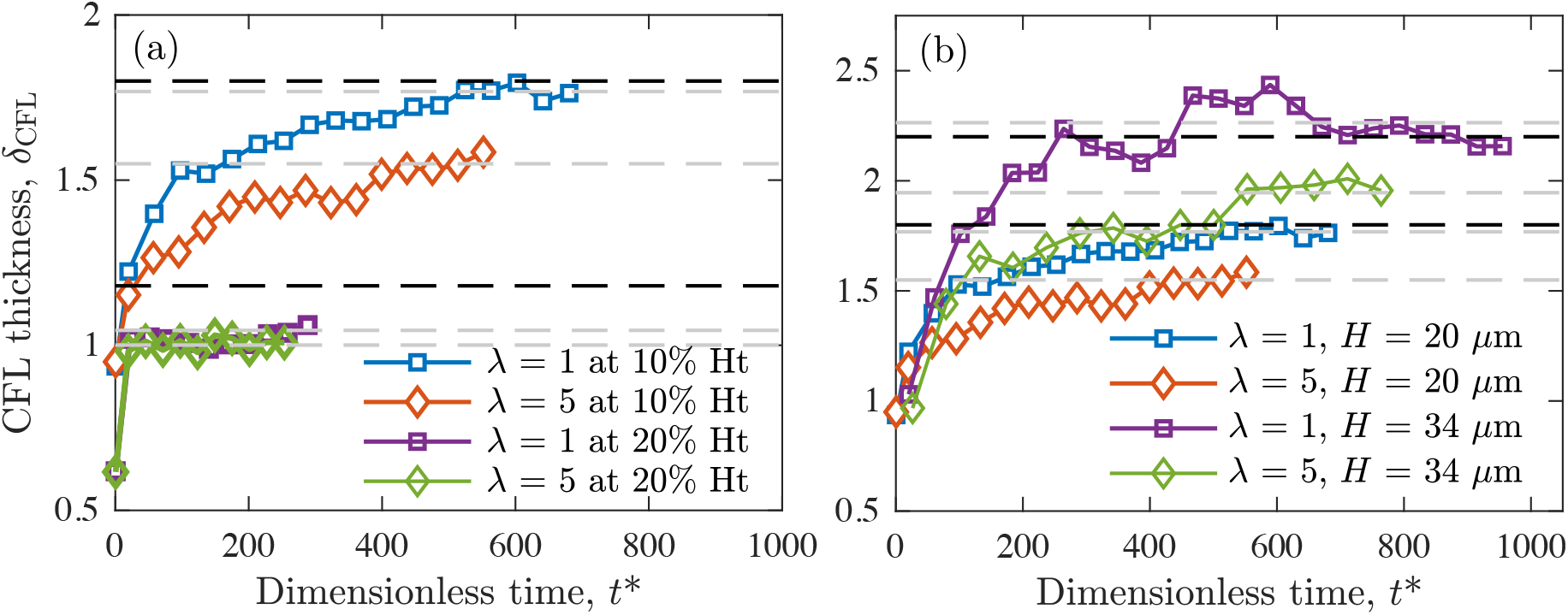
CFL as a function of Ht and λ and time. Additionally the reduced order model results form Qi et al. in 2017 are plotted in dashed lines for λ = 1 as dark dashed lines. In panel (a) we compare the CFL for simulations at two different Ht (10% and 20%) with viscosity ratios of 1 and 5. In panel (b) we examine the effect of channel size and viscosity ratio on the CFL. Note that the CFL is nearly invariant to viscosity ratio for higher volume fraction of 20%. The lower volume fraction of 10% shows good agreement compared to the reduced order model for low viscosity ratio, but the high viscosity ratio deviates significantly from the low λ results from the reduced order model.

In Fig. 4b the CFL as a function of time is plotted for low Ht simulations (10%) at two different confinement ratios (*ε* = 0.28 and *ε* = 0.17). Similar reductions in the CFL size are seen as λ is increased in these simulations and we again see good agreement between the Boltzmann collision model [29] and the steady state CFL for both channel sizes if λ = 1. This allows us to conclude that the results of CFL size reduction are largely generic and only a function of the volume fraction, Ht, and the viscosity ratio, λ.

When examining the lower volume fraction results in Fig. 4b we see that the CFL is influenced by the viscosity ratio (reduced by about 15%). The lowered capacity for lift at the higher viscosity ratio and the nearly constant binary diffusion demonstrated earlier in the single and double particle studies, respectively, lead to this expected result. Additionally, plotted in this figure is the steady, Boltmann collision model results from [30] as dashed lines. We see good agreement between the low viscosity ratio results and the course grained theory. However, it is clear that the effects of viscosity ratio on lift and binary diffusion cause the results to deviate at higher λ and low Ht.

## IV. CONCLUSIONS

In this study we consider the effect of volume fraction (Ht), viscosity ratio (λ), and confinement (*ε*) on cell suspension dynamics. We consider values of the viscosity contrast of 1 and 5 where the latter is more physiologically relevant but the former is utilized considerably in theoretical calculations for simplicity. Additionally, we compare two confinement levels that correspond to appropriate micro-vasculature dimensions (*ε* = 0.28,0.17).

We see that CFL and cell distributions change noticeably when the volume fraction considered is low (Ht = 10%) for all channel sizes. This volume fraction is indeed physiologically relevant for small arterioles. The CFL is reduced in size and the major peaks in the core of the distribution are blunted. We demonstrate for single cells that increasing the viscosity ratio decreases lift but does little to the binary diffusion induced by collision. The weakened driving force towards the center of the channel from lift with the unchanged diffusive contribution likely leads to a reduced CFL size. This result has important implications for simulations of hemostasis in small arterioles since the change in CFL impacts the available space for platelets in physiological conditions.

In addition, we find that at higher volume fractions (Ht = 20%), there are very few changes to the cell distribution and the CFL when the viscosity ratio is changed. In these situations (which are slightly above the physiological hematocrit for arterioles of the size that we are simulating [15, 37]) the excluded volume effects from the large number of cells seem to dominate the behavior leading to small, similarly sized CFLs in both cases.

## ACKNOWLEDGMENTS

The authors of this paper would like to acknowledge support from NSF (CBET-1803765), the US Army High Performance Computation Research Center (AH-PCRC) with the grant number W911NF07200271, and the Open Medicine Foundation (OMF). Computer simulations were performed on AHPCRC’s Excalibur cluster, the Stanford University Certainty cluster, and XSEDE’s COMET cluster.

## Notes

#### Summary of Updates

- Methodology part has shortened and moved to supplementary material. - The Ca number for the lift and binary collision simulation has changed from 1.6 to 1. - The figures corresponding to concentration distribution and the CFL has modified and organized significantly. - The text has been proofread multiple times.

