## Supplementary material for "Effect of Cytoplasmic Viscosity on Red Blood Cell Migration in Small Arteriole-level Confinements": SM: ASaadat_Blood_PRF2019_SM.pdf

Eric S. G. Shaqfeh

*Department of Chemical Engineering,*

*Stanford University, Stanford, CA 94305, USA*

*Department of Mechanical Engineering,*

*Stanford University, Stanford, CA 94305, USA and*

*Institute for Computational and Mathematical Engineering,*

*Stanford University, Stanford, CA 94305, USA*

### I. LIST OF RBC MOVIES

The list of simulation movies and the descriptions are provided in the following table:

TABLE I. List of the movies added as the supplementary material

| Name of the file | Description |
|---|---|
| eps0.28_single_lam1 | A single cell with $\lambda = 1$, Ca=1, and $\epsilon = 0.28$ |
| eps0.28_single_lam5 | A single cell with $\lambda = 5$, Ca=1 and $\epsilon = 0.28$ |
| eps0.28_binary_lam1 | Two cells with $\lambda = 1$, Ca=1 and $\epsilon = 0.28$ |
| eps0.28_binary_lam5 | Two cells with $\lambda = 5$, Ca=1 and $\epsilon = 0.28$ |
| eps0.28_ht10_lam1 | 10% Ht suspension and $\lambda = 1$, Ca=1 and $\epsilon = 0.28$ |
| eps0.28_ht10_lam5 | 10% Ht suspension and $\lambda = 5$, Ca=1 and $\epsilon = 0.28$ |
| eps0.28_ht20_lam1 | 20% Ht suspension and $\lambda = 1$, Ca=1 and $\epsilon = 0.28$ |
| eps0.28_ht20_lam5 | 20% Ht suspension and $\lambda = 5$, Ca=1 and $\epsilon = 0.28$ |

### II. ADDITIONAL METHOD DETAILS

#### A. Numerical Implementation Overview

To solve the coupled fluid-solid problem we utilize an Immersed Finite Element Method (IFEM). More details about this method can be found in a recent publication [5]. To arrive at the governing equations, we rewrite the momentum conservation equations as a single equation over the total domain as follows:

$$\rho \frac{\mathrm{D}\boldsymbol{v}}{\mathrm{D}t} = \nabla \cdot \boldsymbol{\sigma}^{\mathrm{f}} + \boldsymbol{f}^{\mathrm{IB}} \quad \boldsymbol{x} \in \Omega, \tag{1}$$

where $\boldsymbol{f}^{\mathrm{IB}}$ is the immersed boundary force density. It is clear that for conservation of momentum to be satisfied everywhere, the immersed boundary force density must take the following form:

$$\boldsymbol{f}^{\mathrm{IB}} = \nabla \cdot (\boldsymbol{\sigma}^{\mathrm{m}} - \boldsymbol{\sigma}^{\mathrm{f}}) \quad \boldsymbol{x} \in \Omega^{\mathrm{m}}. \tag{2}$$

The discretized immersed boundary method utilizes two separate grids. The Lagrangian grid ($\Omega^{\mathrm{m}}$) while a second fixed Eulerian grid is utilized for the entire domain ($\Omega^{\mathrm{m}} + \Omega^{\mathrm{f}} = \Omega$).

We distinguish between the immersed boundary force on the Lagrangian grid and the immersed boundary force in the Eulerian domain which are defined to be $\boldsymbol{F}^{\mathrm{IB,m}}$ and $\boldsymbol{F}^{\mathrm{IB,f}}$ respectively (note that force densities are given by a lowercase $\boldsymbol{f}$ and forces are given by uppercase $\boldsymbol{F}$).

On the Eulerian domain we therefore solve the following expression with a third order accurate finite volume scheme developed at Stanford's Center for Turbulence research [2]:

$$\rho \frac{\mathrm{D}\boldsymbol{v}}{\mathrm{D}t} = \nabla \cdot \boldsymbol{\sigma}^{\mathrm{f}} + \boldsymbol{f}^{\mathrm{IB,f}} \quad \boldsymbol{x} \in \Omega. \tag{3}$$

We are left to determine the values of $\boldsymbol{F}^{\mathrm{IB,m}}$ for which we utilize finite elements. Details of this expression can be found in a more detailed computational methods paper published elsewhere [5].

Membrane immersed boundary models have additional special considerations. The viscosity of the fluid inside the membrane may not be the same as the exterior fluid. The second important consideration is that the volume of the capsule can drift over time due to interpolation errors, necessitating explicit correction.

For the simulation of RBCs, we solve the following Poisson equation to determine which nodes of the fluid domain are inside the membrane boundaries (this information is encoded as an indicator function $I$):

$$\nabla^2 I = \nabla \cdot \mathbf{G}, \tag{4}$$

where

$$\mathbf{G} = \int_{\Omega^{\mathrm{m}}} \mathbf{n} \mathrm{d}S.$$

For details of this implementation see [1].

Using this information, the RBC internal fluid can be assigned variable viscosity ratios. We can subsequently set the viscosity in the fluid domain to be:

$$\eta_0 = \eta_{\mathrm{out}} + (\eta_{\mathrm{in}} - \eta_{\mathrm{out}}) I.$$

Additionally, since the divergence free character of the flow is not preserved exactly during the velocity interpolation step in the immersed boundary method, the RBCs modeled using a thin membrane may undergo a gradual volume change during the simulation (Note that even though the relative volume change is typically on the order of $10^{-4}$ and smaller in a

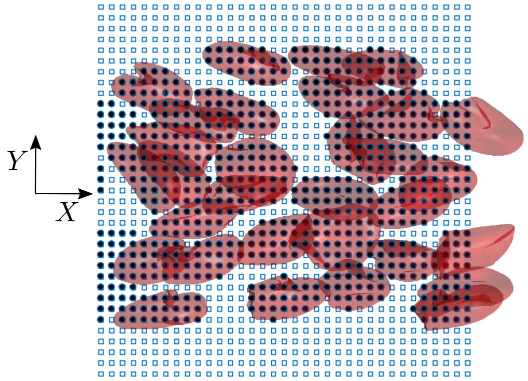


FIG. 1. A graphical representation of how the distribution function is calculated. Points in filled circles are inside the RBCs at this timestep and points marked as open squares are outside (the points represent a discrete sampling of our indicator function, $I$). The integration at each slice in $Y$ gives a more accurate picture of the distribution than the common formation of a histogram from the center of mass locations.

single time step, the associated numerical error will propagate and will cause errors of a few percent by the end of the simulation). In order to avoid this, we exploit the volume conservation algorithm proposed by [3].

##### B. Calculation of Cell Distributions and CFL

The cell distributions will be presented throughout the discussion and we desire to calculate a metric that describes concentration of red blood cells at a given position without having to resort to discrete measures. To calculate the local hematocrit $\mathrm{Ht_L}$, the following integral is calculated over the domain using the indicator function $I$ and the size of the box considered $(L_X, L_Z)$:

$$\mathrm{Ht_L}(Y) = \frac{1}{L_X L_Z} \int \int I(X, Y, Z) dX dZ. \tag{5}$$

The discretized version of this integration can be performed on a regular mesh, and is graphically illustrated in Fig. 1. The points that are inside the RBCs have been illustrated in closed circles and the points outside the RBCs have been illustrated with open squares. Oftentimes, a much coarser measure of the distribution is presented to which typically is

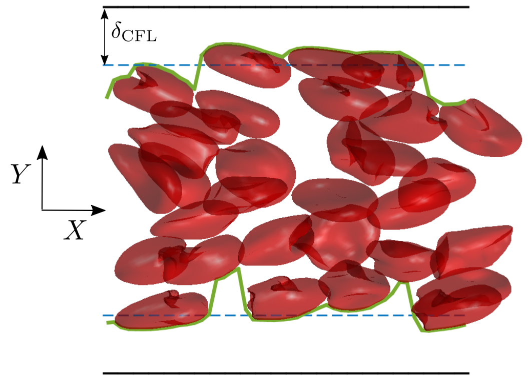


FIG. 2. A graphical representation of how the CFL is calculated. The blue dashed line is the CFL thickness as calculated from the average distance a cell is away from the wall at every $x$-plane. These distances are plotted as a function of $x$ as green solid lines.

a histogram of the center of masses [4, 6]. Utilizing a histogram has the requirement of selecting a sufficiently large bin size to have a smooth variation, but the selected bin size can often qualitatively change how these functions look. However, our approach produces a unique distribution function for a given RBC state for sufficiently small regular mesh size. In this manuscript, the nondimensional size of the mesh is 0.25 in all three directions.

Additionally, we calculate the Cell Free Layer (CFL) thickness for a given RBC state by measuring the distance to the nearest red blood cell from the wall at every $x$ plane and averaging this over the entire domain. This has been illustrated graphically in Fig. 2 where the blue dashed line represents the calculated CFL thickness. The solid green line is the smallest distance from the wall a red blood cell is located in each $x$-plane as a function of $x$. The CFL is calculated as the average distance along this line. To calculate the CFL as a function of time, we use a running average where each data point represents the average in the domain between the two neighboring data points.

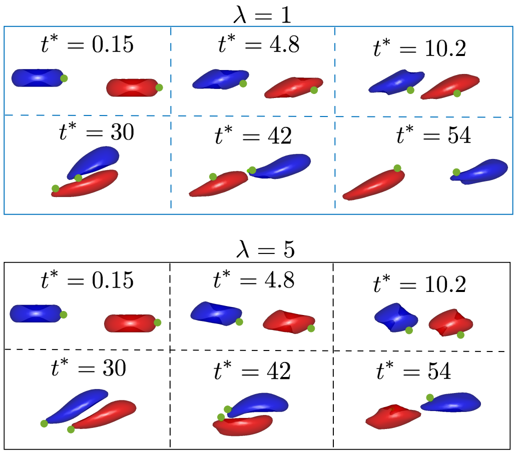


FIG. 3. The collision trajectory at two different viscosity ratios of 1 and 5. The upper row of this figure is at a viscosity ratio of 1 and the lower row is at a viscosity ratio of 5. We note that the higher viscosity ratio exhibits more tumbling motion than the lower viscosity ratio.

#### III. RESULTS AND DISCUSSION

##### A. Binary Cell Diffusion

In Fig. 3 we have presented timelapses images of these two collisions. We note that for the lower viscosity ratio simulations the cells are initially tank treading (the top set of presented images). The higher viscosity ratio collision involves cells that are tumbling instead of tank treading. This likely explains the subtle differences in the collision trajectories presented in the left side of Fig. 3. Ultimately these simple behaviors presented here, lift and binary collision, can be used to construct a collision model.

##### B. Second Moment of Concentration Distribution

In Fig. 4 we have calculated the temporal evolution of the second moment of the concentration profile,

$$M_2 = \sum_Y \left(\bar{N}_{\text{cell}} - N_{\text{cell}}(Y)\right) Y_{\text{com}}^2 \tag{6}$$

where $N_{\text{cell}}(Y)$ is the number of RBCs at a particular height normal to the flow and $\bar{N}_{\text{cell}}$ is its average (the total number of cells normalized by the number of bins.

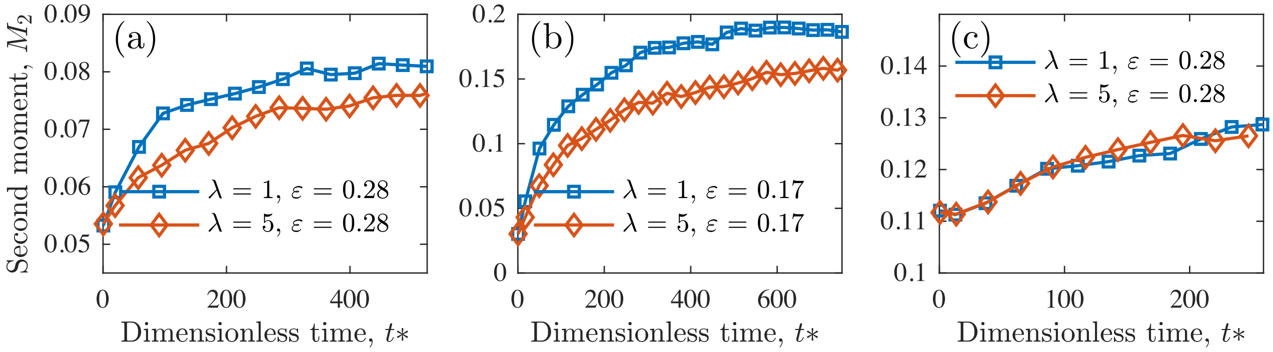


FIG. 4. The second norm of concentration distribution that is presented in Fig. 3 of the manuscript. For the case of $\varepsilon = 0.17$ and 10% Ht, the second moment of $\lambda = 5$ concentration distribution shows up to 30% deviation with respect to the unit viscosity ratio.

---


[1] Bagchi, P. and Kalluri, R. M. (2009). Dynamics of nonspherical capsules in shear flow. *Phys. Rev. E*, 80:016307.

[2] Ham, F., Mattsson, K., and Iaccarino, G. (2006). Accurate and stable finite volume operators for unstructured flow solvers. *Annual Research Briefs*, pages 243–261.

[3] Mendez, S., Gibaud, E., and Nicoud, F. (2014). An unstructured solver for simulations of deformable particles in flows at arbitrary reynolds numbers. *Journal of Computational Physics*, 256:465 – 483.

[4] Qi, Q. (2017). *Understanding Particle Migration, Margination and Adhesion in Cellular Suspensions*. PhD thesis, Stanford University.

[5] Saadat, A., Guido, C. J., Iaccarino, G., and Shaqfeh, E. S. G. (2018). Immersed-finite-element method for deformable particle suspensions in viscous and viscoelastic media. *Phys. Rev. E*, 98:063316.

[6] Zhao, H., Shaqfeh, E. S. G., and Narsimhan, V. (2012). Shear-induced particle migration and margination in a cellular suspension. *Physics of Fluids*, 24(1):011902.
